# An interdependent Cbf1-CCAN interaction stabilizes the budding yeast kinetochore

**DOI:** 10.64898/2026.03.25.714319

**Authors:** Sabrine Hedouin, Changkun Hu, Sue Biggins

## Abstract

Chromosome segregation requires the proper assembly of kinetochores on centromeric DNA. The kinetochore is a complex multi-protein machine comprising more than 40 distinct proteins, but the functional roles of many components remain unclear. One such protein is the yeast transcription factor Cbf1, which directly binds to budding yeast centromeric DNA. Loss of Cbf1 significantly increases the rate of chromosome missegregation, however its precise molecular mechanism of action is unknown. It was recently found that Cbf1 inhibits transcription through the centromere by preventing the untimely pericentromeric transcriptional readthrough via a roadblock mechanism. Intriguingly, restoring the transcriptional roadblock in the absence of Cbf1 binding only partially rescued chromosome missegregation, indicating that Cbf1 performs additional centromeric activities. Here, we show that Cbf1 promotes inner kinetochore assembly both in vitro and in vivo. This assembly function depends on the interface between Cbf1 and Okp1. Moreover, we found that the stable association of Cbf1 with the centromere requires its interaction with the inner kinetochore, revealing an interdependent interaction essential for the assembly and stability of the kinetochore. Thus, Cbf1 functions as a centromere-anchored hub that couples transcriptional roadblocking to CCAN assembly and kinetochore stability.

## INTRODUCTION

Centromeres are essential chromosomal loci required for the accurate partitioning of the genetic material during cell division. They specify the site of assembly of the kinetochore, a conserved multimeric proteinaceous complex that mediates the attachment of chromosomes to dynamic spindle microtubules (1, 2). In most eukaryotes, centromeres are enriched with highly repeated DNA sequences that are poorly conserved. Therefore, most centromeres are epigenetically specified by the incorporation of a centromeric histone H3 variant called centromere protein-A (CENP-A) (3, 4). CENP-A acts as the foundation for the hierarchical recruitment of the inner kinetochore complexes, also called the <u>c</u>onstitutive <u>c</u>entromere-<u>a</u>ssociated <u>n</u>etwork (CCAN) (5). The CCAN then serves as a scaffold for the outer kinetochore that includes the Knl1-Mis12-Ndc80 (KMN or outer kinetochore) network that mediates microtubule attachment and spindle assembly checkpoint signaling (6, 7). Additionally, the CCAN promotes centromere function and maintenance by stabilizing the CENP-A nucleosome, providing additional anchoring on DNA, and contributing to CENP-A replenishment (8–14).

Although there has been tremendous progress in identifying kinetochore components, the specific molecular contributions of many remain unknown. Budding yeast are an excellent model organism to identify their underlying functions because they have one of the simplest kinetochores. Unlike most eukaryotes, the budding yeast centromere is composed of a short ~120 bp unique DNA sequence, called a point centromere, which is both necessary and sufficient for centromere function. It is composed of three conserved centromere DNA elements (CDEI-II-III), which wraps around a single CENP-A^Cse4^ nucleosome (15–17). CDEI is bound by Cbf1, a basal transcription factor with both activator and repressor activity (18, 19). CDEIII is bound by the CBF3 complex, which mediates the deposition of CENP-A^Cse4^ on the AT-rich CDEII element, onto which the rest of the kinetochore assembles (20–22). Interestingly, the stability of the Cse4 nucleosome, the CCAN (also called the Ctf19 complex in budding yeast), and the outer kinetochore are all interdependent (12, 23–26), highlighting the importance of understanding the contribution of each kinetochore component in the assembly and maintenance of the kinetochore.

Cbf1 is one of the first centromere-bound proteins to be discovered in yeast, however, the extent of its roles at the centromere remains elusive. Cbf1 is a homodimeric transcription factor of the basic-helix-loop-helix-leucine zipper family (19). It binds specifically to the pseudo-palindromic sequence CACRTG (R = purine), an E-box motif found at all centromeres and throughout the yeast genome (27–29). Consistent with its broader transcriptional role, *CBF1* deletion leads to methionine auxotrophy and other transcriptional defects (19, 30). Neither Cbf1 nor CDEI is essential, but their deletion results in significant chromosome segregation defects in mitosis and meiosis, indicating that Cbf1 is an important component of the centromere-kinetochore complex. We and others found that Cbf1 negatively regulates centromere transcription (31–33). Its loss leads to constitutive centromere transcription throughout the cell cycle instead of mainly occurring in S-phase. Additionally, by characterizing the centromere transcriptional landscape using Isoform sequencing (Iso-seq), we found that most transcription converging toward the centromere is terminated at the centromere boundaries in a Cbf1-dependent manner, indicating that Cbf1 protects centromeres from transcription via a roadblock mechanism (32). Strikingly, replacing the CDEI element with the binding site of Reb1, a known transcription roadblock factor, was sufficient to block transcription readthrough through the centromere but it only partially rescued the chromosome missegregation defect associated with a mutated CDEI element (32). This indicates that Cbf1 has multiple centromeric functions, with transcriptional regulation only being one of them. Consistent with this possibility, Cbf1 physically interacts with the CBF3 complex and several components of the CCAN (34–36), suggesting it directly participates in kinetochore assembly. However, how these interactions contribute to CCAN assembly and kinetochore stability in vivo remains unknown.

Here we show that Cbf1 functions as a centromere-bound assembly factor that stabilizes the CCAN and, reciprocally, that CCAN assembly enhances Cbf1 retention on centromeric DNA. Cbf1 loss or CDEI mutations reduce the assembly of the Okp1:Ame1 and Ctf19:Mcm21 subcomplexes and almost abrogate the assembly of the Ctf3 and Cnn1:Wip1 subcomplexes. Additionally, we demonstrate that this assembly dependency relies on the binding interface between Cbf1 and the essential CCAN component Okp1. Finally, we found that Cbf1’s stability on the CDEI element is dependent on its multiple interactions with the CCAN, revealing another interdependent interaction required for the maintenance of kinetochore stability and function. Together, our work reveals previously unappreciated feedback between a transcription factor and the inner kinetochore that couples transcriptional roadblocking to kinetochore assembly and stability.

## RESULTS

### Cbf1 is required for CCAN assembly in vitro

The yeast CCAN forms through a hierarchical and interdigitated assembly of multiple subcomplexes (25, 26, 37, 38). Mif2 and Okp1-Ame1, the only essential CCAN components, directly interact with Cse4 and act as the primary recruitment platforms for the outer kinetochore. Ctf19 and Mcm21 associate with Okp1-Ame1 to form the COMA complex that recruits the remaining inner kinetochore components, the Ctf3 and Cnn1 subcomplexes. The Ctf3 complex anchors the CCAN to the DNA via Chl4 and provides an additional accessory linkage to the outer kinetochore through Cnn1-Wip1 (14, 25) (Fig.1A).

**Figure 1.**
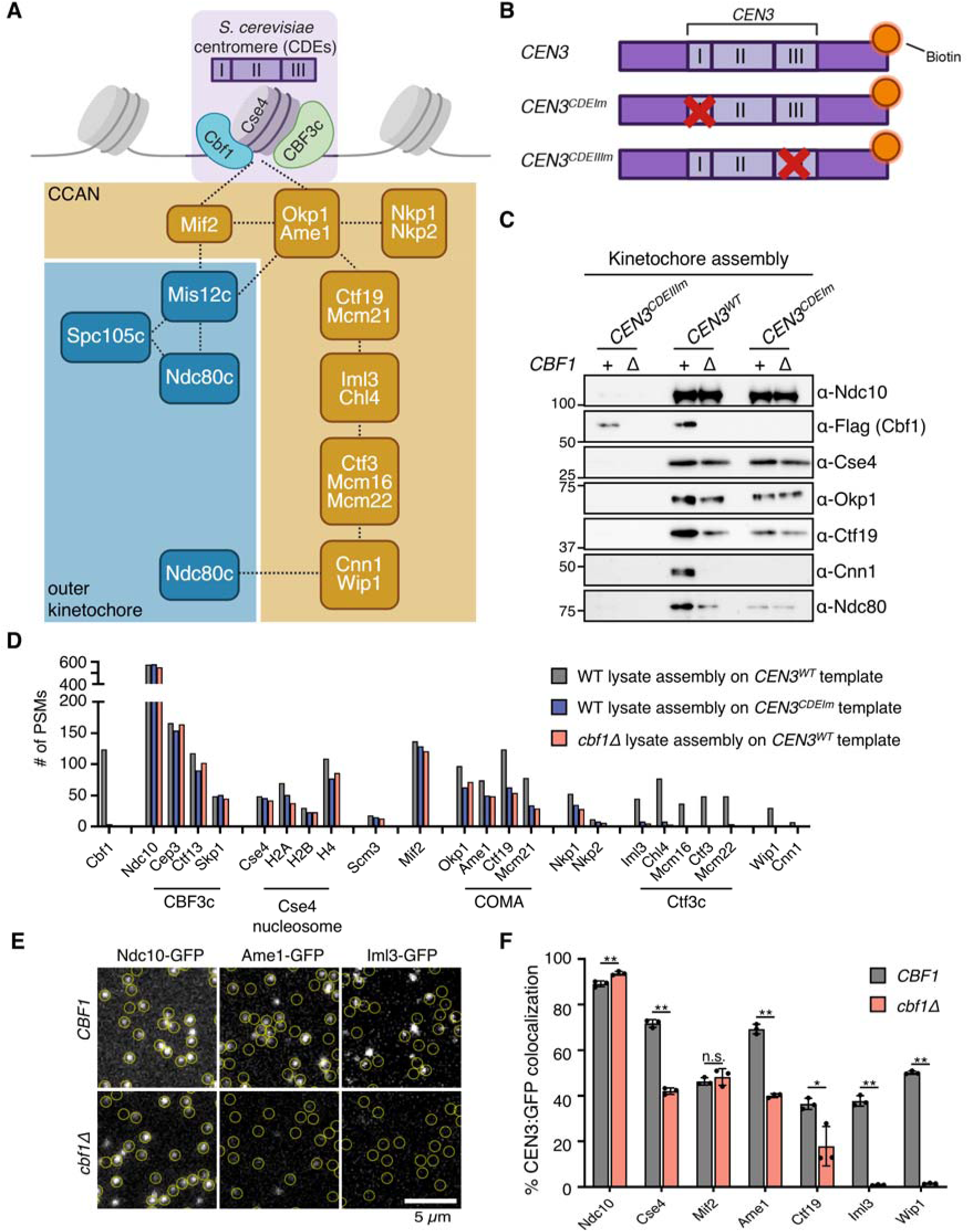
Cbf1 is required for CCAN assembly in vitro. (A) Schematic of the budding yeast centromere and kinetochore. The centromere is composed of three Centromere Determining Elements (CDEs), CDEI, CDEII, and CDEIII, onto which Cbf1, Cse4, and the CBF3 complex assemble, respectively. The CCAN assembles in a hierarchical manner on the Cse4 nucleosome and serves as a platform for outer kinetochore recruitment. CCAN and outer kinetochore subcomplexes are depicted in the yellow and blue boxes, respectively. Dotted lines indicate physical interactions. (B) Schematic of the *CEN3* DNA templates used in the *de novo* in vitro kinetochore assembly assay. The biotinylated DNA templates are depicted in purple, and the red cross indicates mutations in the CDEI or CDEIII elements that disrupt the binding of Cbf1 or CBF3c, respectively. (C) Immunoblots of DNA-bound proteins from de novo kinetochore assembly assays performed using extracts from mitotically arrested WT (*CBF1-3xFLAG* (SBY18421)) and *cbf1*Δ (SBY4958) strains with the indicated DNA templates. (D) Mass spectrometry analysis of DNA bound kinetochore proteins recovered in the kinetochore assembly assay performed as in (B). Raw Peptide-Spectrum Match (PSMs) counts of inner kinetochore proteins are plotted. (E) Representative TIRFM images of surface-tethered *CEN3* DNA molecules (at locations indicated by yellow circles) after incubation for 180 mins with lysates from the following strains: *NDC10-GFP* (SBY22903), *NDC10-GFP cbf1*Δ (SBY24745*), AME1-GFP* (SBY22119), *AME1-GFP cbf1*Δ (SBY24617), *IML3-GFP* (SBY22199), and *IML3-GFP cbf1*Δ (SBY24621). (F) Percentages of colocalization between *CEN3* DNAs and GFP-tagged protein in either in a wild-type or *cbf1*Δ background as analyzed by after 180 min of incubation. Strains used are as follows: *NDC10-GFP* (SBY22903), *NDC10-GFP cbf1*Δ (SBY24745*), CSE4-GFP* (SBY22195), *CSE4-GFP cbf1*Δ (SBY24743), *MIF2-GFP* (SBY22094), *MIF2-GFP cbf1*Δ (SBY24615), *AME1-GFP* (SBY22119), *AME1-GFP cbf1*Δ (SBY24617), *CTF19-GFP* (SBY22116), *CTF19-GFP cbf1*Δ (SBY24619), *IML3-GFP* (SBY22199), *IML3-GFP cbf1*Δ (SBY24621), *WIP1-GFP* (SBY22207), and *WIP1-GFP cbf1*Δ (SBY24625). Error bars represent the standard deviation over three biological repeats. At least 3000 DNA molecules were imaged for each biological replicate. Data for GFP-tagged proteins in a wild-type background were replotted from (23). Statistical significances were analyzed by unpaired t-tests (*, p<0.05; **, p<0.01; n.s., non-significant).

To test whether Cbf1 directly contributes to CCAN localization at the centromere, we used a well-established in vitro kinetochore assembly assay (24), in which kinetochores are *de novo* assembled on a *CEN3* DNA template in budding yeast lysates. We compared lysates from a mitotically arrested *cbf1*Δ strain or a CDEI-mutant CEN DNA template that prevents Cbf1 binding. As a negative control, we used a CDEIII-mutant DNA template that fully abrogates kinetochore assembly (Fig. 1B). Immunoblot analysis of the assembled complexes showed that Ndc10 and Cse4 assembled at levels similar to wild type (Fig. 1C and SI appendix, Fig. S1A). In contrast, the recruitment of several CCAN proteins was either reduced (Okp1 and Ctf19) or absent (Cnn1) when Cbf1 was missing or unable to bind the CDEI-mutant *CEN3* template. This impaired inner kinetochore assembly corresponded with decreased outer kinetochore recruitment, as indicated by reduced Ndc80 levels.

To obtain a more comprehensive view of how Cbf1 loss affects kinetochore assembly, we analyzed the assembled complexes from wild-type, *cbf1*Δ, and CDEI-mutant DNA samples by mass spectrometry and plotted the resulting peptide–spectrum matches (PSMs) counts of inner kinetochore proteins. Consistent with the immunoblot results, most inner kinetochore components, including the CBF3 complex, the Cse4 nucleosome and Mif2 were mostly unchanged in the mutant assemblies (Fig. 1D). Similarly, the COMA complex (Ctf19, Okp1, Mcm21, Ame1) and Nkp1/Nkp2 were roughly reduced by half while the remaining CCAN subunits were completely absent (Fig. 1D).

Next, we employed a recently established single-molecule TIRF microscopy-based kinetochore assembly assay to monitor, with higher sensitivity and kinetic information, the *de novo* recruitment of inner kinetochore proteins on fluorescently labeled *CEN3* DNA in the absence of Cbf1 (12, 23). Briefly, *CEN3* DNAs were tethered to a coverslip and incubated with yeast extract containing GFP-tagged kinetochore proteins for varying durations (up to 180 minutes). The lysate was then washed away, and the slide was imaged to quantify the percentage of DNA molecules colocalizing with GFP. Time-series and end-point imaging largely emulated the trends observed in the bulk kinetochore assembly assay. In the absence of Cbf1, the colocalization of Ame1 and Ctf19 with *CEN3* DNA was reduced by approximately 50%, whereas colocalization of Iml3 and Wip1 was fully abolished (Figs. 1E-F and SI appendix, Fig. S1B). However, we observed one notable difference in the behavior of Cse4, which exhibited reduced stability in the Cbf1-deficient assemblies compared to wild-type. This difference may be explained by the higher sensitivity of the TIRF assay and suggests that Cbf1 either directly stabilizes Cse4 or indirectly stabilizes it through the assembly of Okp1-Ame1 (11, 12). To further quantify these effects, we measured the GFP fluorescence intensity of individual kinetochore proteins colocalizing with DNA after 180 min of assembly. Except for Ndc10, all proteins examined showed reduced fluorescence intensity in the *cbf1*Δ mutant. The magnitude of the reduction ranged from approximately 30% for Cse4 to more than 70% for Ame1 and Ctf19 (SI appendix, Fig. S1C), suggesting reduced occupancy of these proteins within assembled kinetochore particles. Together, our results show that Cbf1 is dispensable for the initial assembly of the CBF3-Cse4-Mif2 centromeric platform but is required for the efficient and complete assembly of CCAN components in vitro.

### CCAN assembly is dependent on Cbf1 in vivo

To determine whether the CCAN’s kinetochore localization depends on Cbf1 in vivo, we purified kinetochores assembled on circular minichromosomes from wild-type or *cbf1*Δ cells (39). Each minichromosome contains a copy of *CEN3*, either wild-type, or mutated in the CDEI or CDEIII elements, and a LacO array to allow purification via LacI-Flag. Immunoprecipitation of the minichromosome via LacI-Flag showed similar trends as the in vitro assemblies: CCAN proteins such as Ctf19 and Cnn1 were largely reduced on CDEI-mutant minichromosomes and on wild-type minichromosomes isolated from the *cbf1*Δ background (Fig. 2A and SI appendix, Fig. S2). We also directly assessed CCAN levels at endogenous centromeres using fluorescence microscopy. Iml3 was tagged at the endogenous locus with mNeonGreen as a representative CCAN protein and visualized together with the spindle pole body marker Spc110-mCherry. We quantified the fluorescence intensity of Iml3-mNeonGreen foci in interphase and mitotic cells and, as expected from the assembly assays, found that it was significantly reduced in *cbf1*Δ cells (Figs. 2B and 2C).

**Figure 2.**
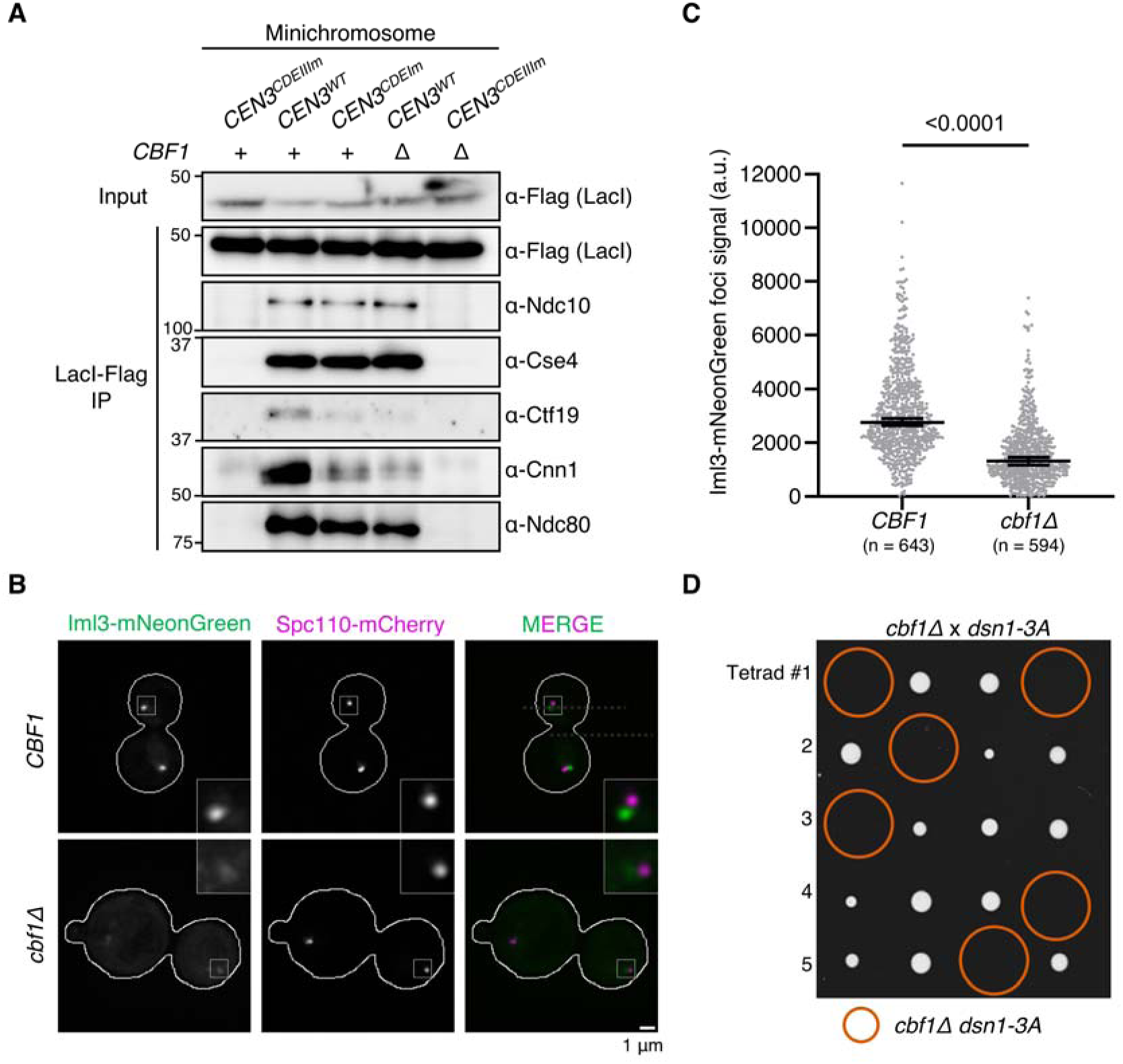
Cbf1 is required for CCAN assembly in vivo. (A) Immunoblot of kinetochore proteins associated with purified minichromosomes. *CEN3-LacO* plasmids carrying the indicated mutations were purified via LacI-Flag immunoprecipitation from either wild type or *cbf1*Δ cells. Minichromosome-bound kinetochore proteins were analyzed by immunoblotting with the indicated antibodies. Strains used were: *CEN3^CDEIIIm^-LacO* (SBY19146), *CEN3-LacO* (SBY19145), *CEN3^CDEIm^-LacO* (SBY19147), *CEN3-LacO cbf1*Δ (SBY19149), and *CEN3^CDEIIIm^-LacO cbf1*Δ (SBY19150). (B) Fluorescence microscopy of Iml3-mNeonGreen and Spc110-mCherry in wild-type (SBY24879) or *cbf1*Δ (SBY24881) asynchronously growing cells. Representative images of maximum intensity projection of single cells are shown. Insets are magnified an additional 3X. Scale bar, 1 μm. (C) Quantification of background-corrected Iml3-mNeonGreen fluorescence intensity at kinetochore foci from (B). Horizontal lines indicate the median and error bars represent the 95% confidence interval of two independent experiments. The number (n) of total kinetochore foci analyzed is noted on the graph. Statistical significance was assessed using a two-tailed Mann–Whitney U test. (D) Tetrad dissection of a cross between *cbf1*Δ (SBY4958) and *dsn1-3A* (SBY14171) strains. The four spores from individual asci are aligned in horizontal rows. Orange circles represent spores with double mutant genotype.

Although most CCAN components are non-essential in budding yeast, they exhibit synthetic lethality when combined with the *dsn1-3A* allele, a phospho-null allele of Dsn1 that downregulates the recruitment of the outer kinetochore (24, 40). We therefore hypothesized that if Cbf1 promotes CCAN recruitment, its deletion should be synthetically lethal with the *dsn1-3A* allele, which was indeed the case (Fig. 2D). Together, our data show that Cbf1 is required for complete CCAN localization at the endogenous yeast centromere.

### Cbf1 recruits the CCAN separately from its transcription roadblock activity

We next tested whether the Cbf1 roadblock activity is involved in CCAN kinetochore localization. We previously found that replacing CDEI with a Reb1 binding site (*CEN3^CDEI::Reb1-BS^*), restores the transcriptional roadblock in vivo (32). We therefore immunoprecipitated kinetochores assembled on a minichromosome containing the chimeric *CEN3^CDEI::Reb1-BS^* (Fig. 3A). We found that the mutant minichromosome lacked CCAN binding similarly to those purified from the *CEN3^CDEIm^* minichromosome (Fig. 3B and SI appendix, Fig. S3A), strongly suggesting that the role of Cbf1 in localizing the CCAN to the centromere is separate from its transcription roadblock activity.

**Figure 3.**
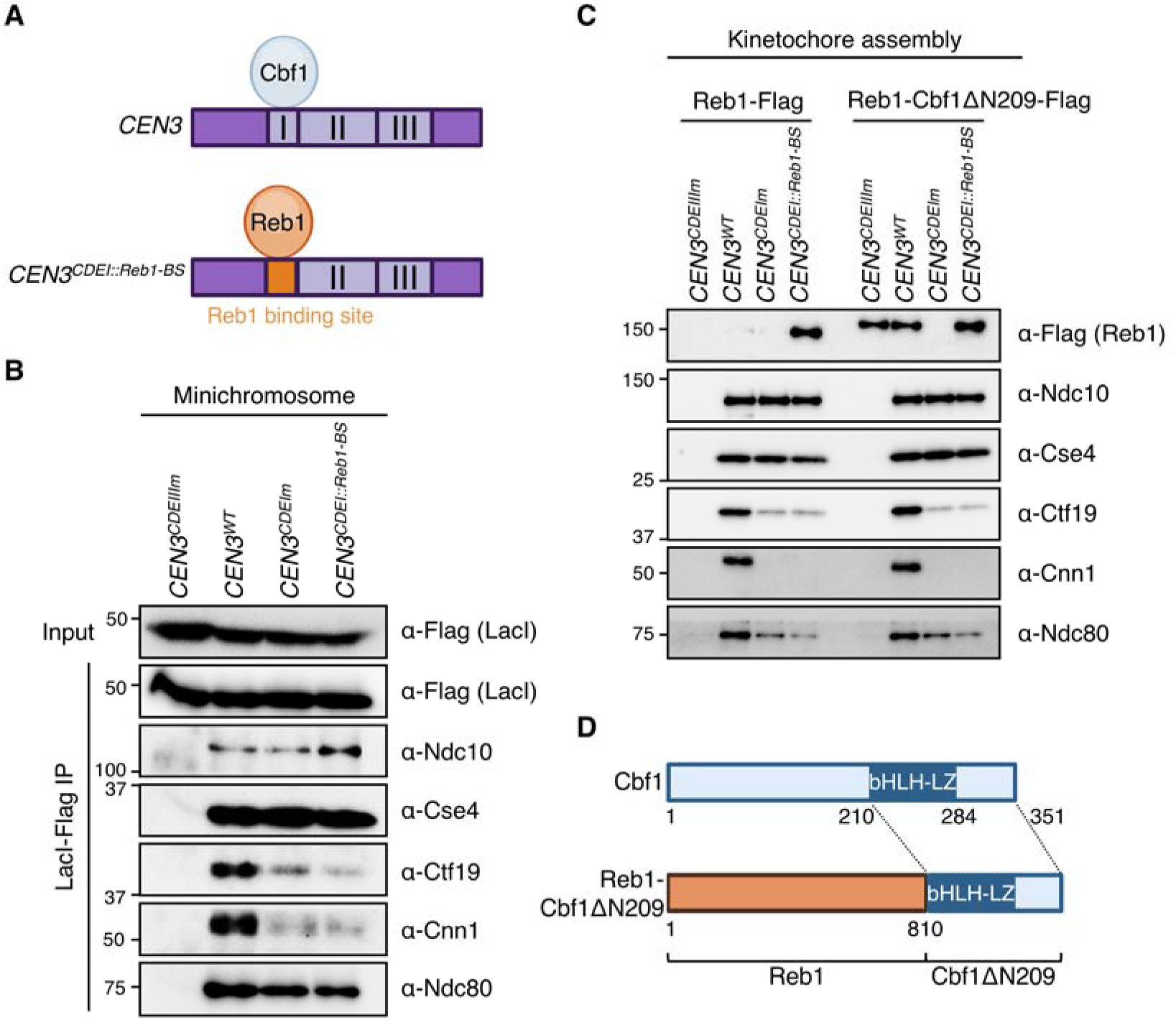
Cbf1’s role in CCAN recruitment is separate from its transcriptional roadblock activity. (A) Schematic of the wild-type *CEN3* template which binds Cbf1 at the CDEI element, and the *CEN3^CDEI::Reb1-BS^* template where the CDEI element is replaced by the Reb1 binding site to ectopically recruit Reb1 to the centromere. (B) Immunoblot of kinetochore proteins associated with purified minichromosomes. *CEN3-LacO* plasmids carrying the indicated mutations were purified via LacI-Flag immunoprecipitation from otherwise wild-type cells. Minichromosome-bound kinetochore proteins were analyzed by immunoblotting with the indicated antibodies. Strains used are as follows: *CEN3^CDEIIIm^-LacO* (SBY19146), *CEN3-LacO* (SBY19145), *CEN3^CDEIm^-LacO* (SBY19147), and *CEN3^CDEI::Reb1-BS^-LacO* (SBY19148). (C) Immunoblots of DNA-bound proteins from de novo kinetochore assembly assays performed using extracts from asynchronously grown *REB1-3xFLAG* (SBY25315) and *REB1-cbf1*Δ*N209-3FLAG* (SBY25317) strains with the indicated DNA templates. (D) Schematics of Cbf1’s domain architecture and of the chimeric Reb1-Cbf1ΔN209 protein. Reb1-Cbf1ΔN209 corresponds to full length Reb1p fused at the C-terminus with a truncated Cbf1 starting at amino acid 210 and comprising the bHLH-LZ and C-terminal domains.

We next asked whether Cbf1 needs to directly bind to the CDEI element to localize the CCAN. To test this, we performed a kinetochore assembly assay on the *CEN3^CDEI::Reb1-BS^*template. First, we tested whether Reb1 binding could restore CCAN localization in vitro. As expected, Reb1 bound to the *CEN3* DNA solely when its binding site was present (Fig. 3C and SI appendix, Fig. S3B). Both Ndc10 and Cse4 were recruited to the *CEN3^CDEI::Reb1-BS^* template normally. However, Ctf19, Cnn1 and Ndc80 failed to assemble properly, to a similar extent as assemblies on a CDEI-mutant template, consistent with Reb1 binding not restoring CCAN localization to minichromosomes in vivo. Next, we tested whether the ectopic recruitment of Cbf1 to the kinetochore could rescue CCAN assembly. We fused the C-terminal domain of Cbf1, containing the basic helix-loop-helix-leucine zipper (bHLH-LZ) domain that is sufficient for ensuring both normal chromosome segregation and transcriptional regulation activities (28), to the endogenous Reb1 protein (Reb1-Cbf1ΔN209-Flag) (Fig. 3D). Expression of the fusion protein did not impair cell viability, indicating that the essential functions of Reb1 remained intact (SI appendix, Fig. S3C). We then performed an assembly assay and found that the fusion protein associated with *CEN3* DNA either through its Cbf1 moiety when CDEI was intact, or through its Reb1 moiety on the chimeric *CEN3^CDEI::Reb1-BS^* DNA (Fig. 3C and SI appendix, Fig. S3B). Expression of the Reb1-Cbf1ΔN209-Flag protein did not alter kinetochore assembly on wild-type DNA. However, it failed to rescue CCAN localization on the *CEN3^CDEI::Reb1-BS^* template (Fig. 3C and SI appendix, Fig. S3B), suggesting that Cbf1’s kinetochore assembly activity requires its direct DNA binding or that the fusion protein does not adopt a conformation compatible with CCAN interaction. Together, these data indicate that Cbf1 promotes CCAN localization through a mechanism that requires the protein to bind to CEN DNA and that is separable from its transcription roadblock activity.

### Cbf1 and CCAN stability at the centromere are interdependent

We next sought to determine how Cbf1 stabilizes the CCAN at the centromere. A recent structural study of a reconstituted CCAN complex identified key interactions between Cbf1 and several CCAN components including Okp1, Ctf19, Chl4, Iml3 and Cnn1/Wip1 (35). Mutations predicted to disrupt the Cbf1-Okp1 interaction sensitized cells to the microtubule depolymerizing drug benomyl to a similar extent as observed for *cbf1*Δ cells (35). Therefore, we hypothesized that this interaction might be essential for the stabilization of the CCAN. To test this, we mutated the same residues on Cbf1, L283E and L287W, which are involved in the Cbf1-Okp1 interface (Cbf1-EW) (Fig. 4A) (35). Immunoprecipitation of Cbf1 followed by immunoblotting against Okp1 validated a weaker association between Cbf1-EW and Okp1 *in vivo* (Fig. 4B). We next assayed lysates made from cells containing the Cbf1-EW mutant in the in vitro kinetochore assembly assay. Cbf1-EW maintained its ability to bind to the CEN DNA, albeit to a slightly lower degree than wild-type Cbf1, suggesting it may be partially unstable (Fig. 4C and SI appendix, Fig. S4A). Interestingly, CCAN assembly in the *cbf1-EW* extracts was impaired to a similar extent as observed when using *cbf1*Δ extracts, indicating that the Cbf1-Okp1 interface is critical for CCAN localization to the centromere (Fig. 4C and SI appendix, Fig. S4A). To confirm the involvement of the Cbf1-Okp1 interaction in CCAN function in vivo, we crossed the *cbf1-EW* mutant to the *dsn1-3A* mutant and found that the double mutants are synthetically lethal (Fig. 4D).

**Figure 4.**
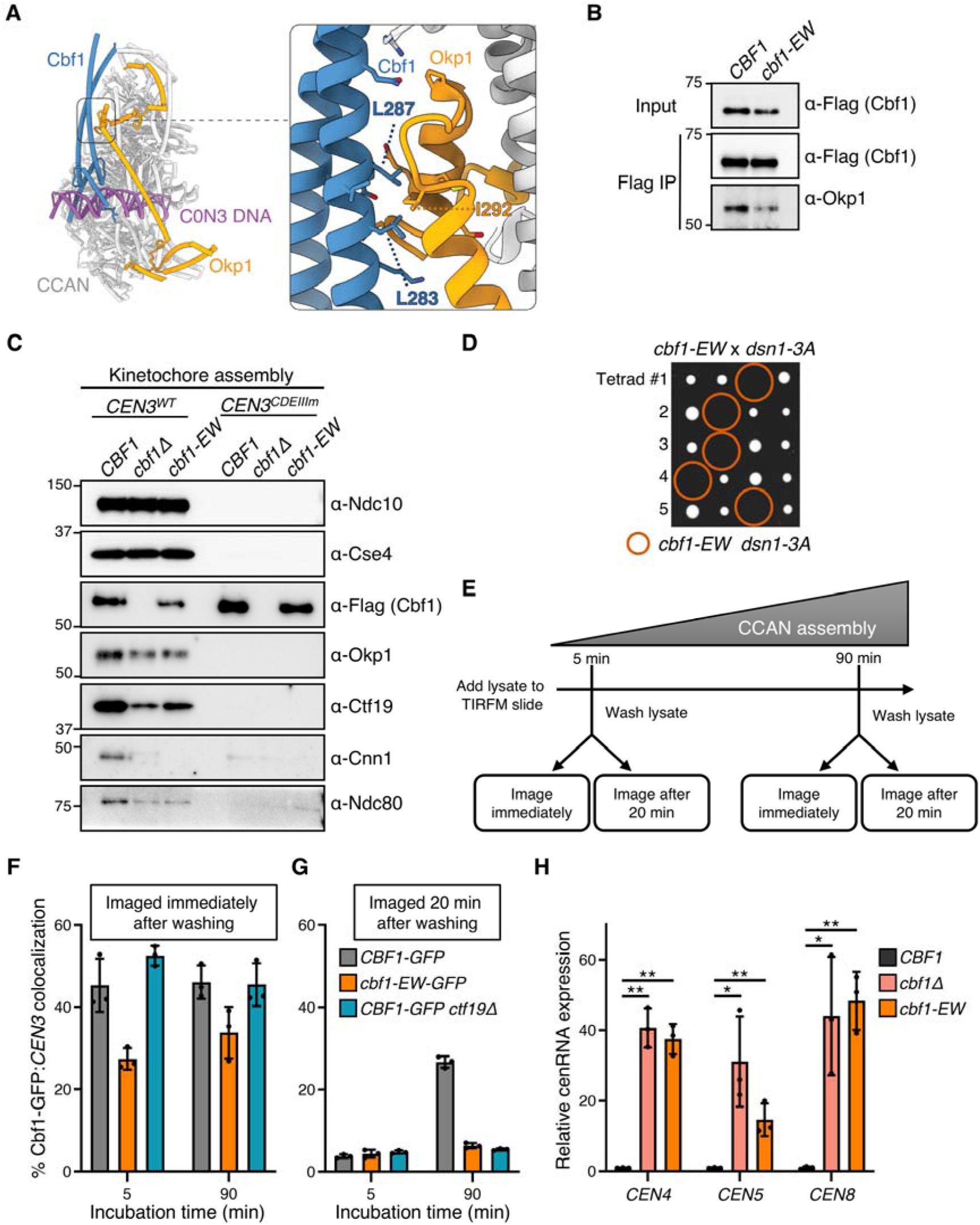
Interdependent stability of Cbf1 and CCAN at the centromere. (A) Structure of the Cbf1:CCAN complex bound to CDEI DNA (PDB: 8OVW). The interaction between Cbf1 and Okp1 is detailed in the inset. (B) Flag-tag immunoprecipitations of Cbf1-3Flag (SBY18421) and Cbf1-EW-3Flag (L283E, L287W; SBY22227) were immunoblotted against Okp1 to analyze co-purifying levels. (C) Immunoblots of DNA-bound proteins from de novo kinetochore assembly assays performed using extracts from asynchronously grown *CBF1-3FLAG* (SBY18421), *cbf1*Δ (SBY4958), and *cbf1-EW-3FLAG* (SBY22227) strains with the indicated DNA templates. (D) Tetrad dissection of a cross between *cbf1-EW* (SBY22227) and *dsn1-3A* (SBY14171) strains. The four spores from individual asci are aligned in horizontal rows. Orange circles represent spores with double mutant genotype. (E) Schematic of the TIRFM stability assay. Lysate is incubated on the TIRFM slide for 5 or 90 minutes before being washed off. Slides were then either imaged immediately or after 20 minutes. (F) Percentages of colocalization between *CEN3* DNAs and Cbf1-GFP (SBY22129), Cbf1-EW-GFP (SBY22923) and Cbf1-GFP in *ctf19*Δ cells (SBY24889) as analyzed by TIRFM after 5 min or 90 min of incubation and imaged immediately post wash. Error bars represent the standard deviation over three biological repeats. At least 3000 DNA molecules were imaged for each biological replicate. (G) Same as in (E) but imaged 20 minutes post wash. (H) RT-qPCR analysis of cenRNA expression of *CEN4*, *CEN5*, and *CEN8* in wild type (SBY22452), *cbf1*Δ (SBY22454), and *cbf1-*EW (SBY22456) cells arrested in G1 with α-Factor. Expression levels were quantified relative to that of wild type (mean ± SD, n=3). Statistical significances were analyzed by unpaired t-tests (*, p<0.05; **, p<0.01).

Recent work from our lab shed light on the interdependency of yeast kinetochore subcomplexes for their centromere association (23). Surprisingly, we found that outer kinetochore components, such as the Mtw1 complex, are required for the stability of inner kinetochore proteins (23). We therefore wondered whether the reduced association of the Cbf1-EW mutant on centromeres in the in vitro kinetochore assembly might reflect a role for CCAN stabilizing Cbf1 on the CDEI element. To test this, we utilized the single molecule TIRF microscopy assay to directly assess Cbf1 localization on CEN DNA either when its interaction with Okp1 is compromised or in a non-essential CCAN mutant that disrupts most CCAN assembly except for Okp1-Ame1 (*ctf19*Δ) (24, 26). We flowed Cbf1-GFP lysates onto slides and incubated for either 5 or 90 minutes, washed off the lysate, and immediately monitored Cbf1 colocalization with fluorescently labeled *CEN3* DNA (Fig. 4E). Wild-type Cbf1 was rapidly assembled on the DNA, within 5 minutes of lysate incubation, at roughly 45% colocalization, and persisted at similar levels after 90 minutes of incubation (Fig. 4F). In contrast, less Cbf1-EW associated with the DNA than wild-type Cbf1 at both incubation time points, consistent with the bulk assembly assay. However, Cbf1 binding was not affected in a *ctf19*Δ background that does not alter Okp1 levels, consistent with Okp1 likely being the major Cbf1 interactor that promotes CCAN assembly (Fig. 4F). Thus, Cbf1’s centromere localization is partially dependent on its interaction with Okp1.

Next, we evaluated whether Cbf1’s long-term stability on DNA is dependent on the CCAN. To test this, we used the same TIRFM assay described above but we instead monitored how much of the Cbf1-GFP-*CEN3* colocalization was retained 20 minutes after the lysate was washed off (Fig. 4E). We found that early assembled wild-type Cbf1 was highly unstable, with colocalization dropping from 45% immediately after wash-off to 5% after 20 minutes (Figs. 4F-G, 5-minute incubation time). Conversely, Cbf1 from wild-type lysates remained more stable at the 90-minute time point, with levels ranging from 45% right after wash-off to 25% after 20 minutes (Fig 4F-G, 90-minute incubation time). These data suggest that although Cbf1 binds rapidly to the CDEI element, it is initially unstable and becomes progressively more stable over time. In line with this, we recently found that the CCAN assembles gradually in the TIRFM assay, increasing from less than 10% assembly at 5 minutes to roughly 80% by 90 minutes (23). Together, these findings indicate that the enhanced stability of Cbf1 on DNA correlates with the progressive recruitment of the CCAN. Strikingly, neither Cbf1-EW nor Cbf1 from a *ctf19*Δ lysate were stable on the DNA at either time point (Fig. 4G), consistent with the idea that Cbf1 stability on the CDEI element is dependent on multiple CCAN interactions. We therefore suspected that the Cbf1-EW mutant would also be compromised in its centromere transcription roadblock activity because of its instability and indeed we found it had increased cenRNA transcription levels (Fig. 4H). This transcriptional effect is likely localized to the centromere as *cbf1-EW* cells did not exhibit the methionine auxotrophy that exists in *cbf1*Δ cells (SI appendix, Fig. S4B). Taken together, these results provide new insights into the interdependency between Cbf1 and the CCAN for their stability at the yeast centromere.

## DISCUSSION

Here we identified a previously unknown function of Cbf1 in the regulation of kinetochore assembly. In contrast to prior models in which transcription factors at centromeres primarily regulate transcription, our data uncover a feedback module in which a DNA-binding factor and the inner kinetochore mutually reinforce each other to build a robust kinetochore. In accordance with recent structural studies that identified multiple interactions between Cbf1 and CCAN subunits, we found that Cbf1 is directly required for the stable and integral assembly of the CCAN, which in return stabilizes the overall kinetochore assembly. We tested whether blocking centromeric transcription was connected to CCAN localization by tethering Reb1 to the centromere in lieu of Cbf1. Although Reb1 restores roadblock activity, it did not restore CCAN localization. Therefore, the Cbf1-CCAN interaction is critical for CCAN localization and is a distinct activity from Cbf1’s roadblock activity.

Interestingly, our data show that Cbf1 loss affects the assembly of CCAN subunits to different degrees, with the COMA and Nkp1/Nkp2 complexes being reduced while the Ctf3 complex is almost absent. Given the hierarchical assembly of the CCAN on CEN DNA (Fig. 1A) (23–25), first Okp1-Ame1 followed by the rest of the CCAN, any perturbation of Okp1-Ame1 stability is likely to impede the assembly of downstream subcomplexes. Consistent with this, the reduced interaction between Cbf1-EW and Okp1 disrupts Ctf3 complex assembly to a similar extent as a *cbf1*Δ mutant in the in vitro kinetochore assembly assay. Cbf1 also contacts the COMA and Ctf3 subcomplexes, indicating that each interaction might cooperatively strengthen the overall stability of the complete CCAN. While a reconstituted CCAN forms a stable complex in vitro in the absence of Cbf1 (14, 41, 42), our results show the clear dependency of native CCAN on Cbf1 for its full assembly on centromeric DNA. Future studies will be needed to determine whether Cbf1 promotes CCAN assembly, retention, or both, and to assess the contribution of its additional interactions with CCAN components to the kinetochore assembly process.

Our TIRFM data show that Cbf1’s stability at the centromere increases correlatively with CCAN arrival and diminishes when Cbf1-CCAN interactions are impaired. Therefore, although Cbf1 directly recognizes and binds its CDEI DNA motif, its kinetochore retention is dependent on its interaction with the whole CCAN. This result sheds light on an earlier study that showed discrepancies of Cbf1 binding affinity between in vitro and in vivo experiments and proposed that some unknown kinetochore proteins help stabilize Cbf1 in vivo (43). Thus, our work identifies Okp1 as a key partner for Cbf1 stability on CEN DNA and helps explain why Cbf1 is more stable at the kinetochore in vivo. Furthermore, we recently discovered multiple interdependencies between inner and outer kinetochore subunits governing their centromere localization (12, 23), suggesting that overall kinetochore stability in vivo requires many complex protein interactions.

Cbf1 exhibits additional direct interactions with kinetochore proteins, besides the CCAN, which may also stabilize itself on DNA and/or the kinetochore. Notably, Cbf1 was shown to interact directly with Ndc10, Cep3 and Ctf13 of the CBF3 complex in vitro and both Cep3 and Skp1 can, individually or in combination, stimulate Cbf1’s DNA binding activity (34, 36). Moreover, Cbf1 binding to CDEI is dependent on the CBF3 complex in vivo (44) and multiple negative genetic interactions between Cbf1 and CBF3 complex members have been reported (45, 46). However, our discovery that the CCAN stabilizes Cbf1 allows for another interpretation of these results. Given that the CBF3 complex promotes Cse4 nucleosome deposition and hence CCAN recruitment, it is possible that the effect observed on Cbf1 stability is indirect and the result of CCAN assembly defects. Nonetheless, determining if a direct Cbf1-CBF3 interaction promotes Cse4 nucleosome stability (34) or Cbf1-CCAN stability is an interesting open question that will require separation of function alleles that are difficult to make because Cbf1 and Ndc10 interact via their DNA-binding domains (34).

It was previously found that Cbf1 can bend centromeric DNA (47), which could influence centromeric chromatin architecture and thereby affect the assembly or stability of the Cse4 nucleosome and associated kinetochore proteins. However, several observations argue that DNA bending alone is unlikely to account for the function of Cbf1. Previous work showed that replacement of CDEI with an artificial intrinsic DNA bend does not rescue the chromosome segregation defects associated with loss of Cbf1 (47), indicating that the presence of the protein itself is required. Consistent with this idea, recruitment of Reb1, a DNA-binding protein that induces a similar DNA bend (48), failed to restore CCAN assembly in our assays. Together, these findings suggest that the contribution of Cbf1 to kinetochore assembly cannot be explained solely by its effects on DNA architecture and is more likely to involve specific interactions with CCAN components.

Although both transcription roadblock and kinetochore stability are distinct Cbf1 activities, they are likely inseparable and require stable and direct binding to the CDEI element (49). Indeed, the ectopic recruitment of Cbf1 to the centromere via a Reb1 fusion failed to rescue CCAN assembly, suggesting that the conformation adopted by the fusion protein was unable to make proper Cbf1-CCAN contacts. Likewise, reduced Cbf1-CCAN interaction destabilized Cbf1 on DNA, contributing to the loss of CEN transcriptional repression which further suggests that the CCAN participates, directly or indirectly, in the transcriptional roadblock at centromeres. Altogether, our work reveals a feedback module whereby a DNA-binding factor and the inner kinetochore mutually stabilize each other to establish a robust kinetochore, paralleling the cooperative role of CENP-B with CENP-A and CCAN components in maintaining centromere integrity in mammals (50–53). In the future, it will be interesting to determine whether DNA-binding proteins that both regulate centromeric transcription and stabilize CCAN assembly represent an evolutionary conserved design principle of centromere biology.

## MATERIAL AND METHODS

### Strain construction and microbial techniques

The *Saccharomyces cerevisiae* strains used in this study are listed in SI appendix, Table S1 and are derived from SBY3 (W303). Standard genetic crosses, media and microbial techniques were used (54). Gene deletions and epitope tagged alleles were constructed at the endogenous loci by standard PCR-based integration as described in (55) and confirmed by PCR. Plasmid mutagenesis was performed as described (56). The plasmids and primers used to generate strains are listed in SI appendix, Tables S2 and S3, respectively. All liquid cultures were grown in yeast peptone dextrose rich (YPD) media supplemented with 2% glucose (Sigma-Aldrich) and 0.02% adenine (MP Biomedicals) at 23 °C unless noted otherwise. For minichromosome immunoprecipitations, cells were grown in YC-TRP supplemented with 2% glucose and 0.02% adenine to maintain selection for the minichromosome. For microscopy, yeasts were grown in synthetic complete (SC) media, supplemented with 2% glucose and 0.06% adenine.

### *De novo* kinetochore assembly assay

Preparation of DNA templates for the kinetochore assembly assay were performed as described in (24). Plasmids and primers used to generate the DNA templates are listed in SI appendix, Tables S2 and S3, respectively. M280 streptavidin Dynabeads (Invitrogen, #11205D) were coated with biotinylated *CEN3* DNA. *De novo* kinetochore assembly assay was performed with yeast whole cell extract as described in (24). Briefly, 250 μL of whole cell extract and 10 μL of DNA-coated M280 Dynabeads were incubated at room temperature for 90 min to allow kinetochore assembly. Beads were then washed 4 times with 1 mL of Buffer L (25 mM HEPES pH 7.6, 2 mM MgCl2, 0.1 mM EDTA pH 7.6, 0.5 mM EGTA pH 7.6, 0.1% NP-40, 175 mM K-Glutamate, and 15% Glycerol) supplemented with protease inhibitors (10 mg/ml leupeptin, 10 mg/ml pepstatin, 10 mg/ml chymostatin, 0.2 mM PMSF) and 2 mM DTT. Bound proteins were eluted by resuspending the beads in 20 μL of SDS buffer, boiling the beads at 100 LJC for 3 min, and collecting the supernatant. Bound proteins were examined by immunoblotting, described below. All experiments were repeated two or more times as biological replicates to verify reproducibility, and a representative result is reported.

### Mass spectrometry

Following the standard kinetochore assembly protocol and washes, assembled kinetochores were washed twice with 1 mL of 50 mM HEPES pH 8, then resuspended in ~60 μL of 0.2% RapiGest SF Surfactant (Waters) in 50 mM HEPES pH 8. Proteins were eluted by gentle agitation at room temperature for 30 min. A small portion of the eluate was added to SDS buffer and analyzed by SDS-PAGE and immunoblotting. The remaining sample was sent to the Fred Hutchinson Cancer Center Proteomics & Metabolomics shared resource facility for LC/MS/MS analysis as described in (57). Mass spectrometry data generated in this study are available through Mass Spectrometry Interactive Virtual Environment (MassIVE, University of California San Diego) located here: https://massive.ucsd.edu/ProteoSAFe/dataset.jsp?task=22882ab4ea494f0288c94dfd7b2dc408.

### Protein and minichromosome immunoprecipitation

Protein and minichromosome immunoprecipitations were performed as previously described (39, 58). Briefly, cells were grown until OD_600_ ≈ 1.2 for minichromosome immunoprecipitation or OD_600_ ≈ 3 for protein immunoprecipitation, harvested, resuspended in Buffer H/0.15 (25□mM Hepes, pH 8.0, 2□mM MgCl_2_, 0.1□mM EDTA, 0.5□mM EGTA, 0.1% NP-40, 15% glycerol, and 150□mM KCl) containing phosphatase inhibitors (0.1□mM Na-orthovanadate, 0.2 µM microcystin, 2□mM β-glycerophosphate, 1□mM Na pyrophosphate, and 5□mM NaF) and protease inhibitors (20 µg/ml leupeptin, 20 µg/ml pepstatin A, 20 µg/ml chymostatin, and 200 µM PMSF) and then flash frozen. Cells were lysed in a freezer mill (SPEX SamplePrep) and then centrifuged at 13,000□g for 30 minutes at 4 °C. Lysate protein concentration was measured using Pierce BCA Protein Assay Kit (Thermo Scientific) and equivalent amounts of input protein were utilized for each experiment. Protein G Dynabeads (Invitrogen) were crosslinked with an α-M3DK antibody that recognizes the 3Flag epitope tag (59) (Genscript) and immunoprecipitations were performed at 4 °C for 3 hours. Beads were washed twice with lysis buffer containing 2□mM DTT and protease inhibitors, three times with lysis buffer with protease inhibitors, and proteins were eluted by gentle agitation of beads in elution buffer (Buffer H + 0.5□mg/ml 3FLAG Peptide [Sigma-Aldrich]) for 30□min at room temperature.

### Immunoblotting

Proteins eluted from a kinetochore assembly assay or immunoprecipitation were separated on a 10% SDS-PAGE gel. Proteins were transferred to a 0.45 μm nitrocellulose membrane (BioRad) and standard immunoblotting was performed. Primary antibodies were used as follow: α-Ndc10 (OD1) 1:5,000 and α-Ndc80 (OD4) 1:5,000 were a generous gift from Arshad Desai; α-Cse4 (9536) 1:500; α-Ctf19 (498) 1:5,000; α-Cnn1 1:1,000; α-Okp1 (4128) 1:5,000; α-M3DK (anti-Flag, Genscript) 1:3,000. HRP conjugated secondary antibodies (GE Healthcare) were detected with Pierce SuperSignal West Dura enhanced chemiluminescent (ECL) substrate (ThermoFisher Scientific, #PI-34076). Immunoblots were imaged with a ChemiDoc MP system (Bio-Rad).

### Fluorescence microscopy and quantification

Log-phase cultures were grown in synthetic complete (SC) media (supplemented with 2% glucose and 0.06% adenine). 1 mL of cells was harvested at 3,000 x g for 2 min and resuspended in 50 μL SC media supplemented with 2% glucose and 0.06% adenine. 2.5 μL of cells were mounted onto a 1.5% agarose pad (dissolved in SC media) assembled onto a microscope slide and sealed with liquid VALAP (33% petroleum jelly, 33% lanolin, 33% paraffin) prior to imaging. Images were acquired on a DeltaVision Ultra using a 100x/1.4 PlanApo N oil immersion objective (Olympus) using a 16-bit scientific complementary metal-oxide semiconductor detector at 22 °C. Z-stacks of cells were obtained at 0.2 μm sections. Images were deconvolved in softWorX v7.2.1 (GE) using standard settings. Projections were made using a maximum-intensity algorithm.

For Iml3-3mNeonGreen fluorescence intensity quantification, using maximum intensity projections, kinetochores from both interphase and mitotic cells were circled using the freehand tool in FIJI (ImageJ, v1.54p) and the integrated fluorescence intensity was measured and background-corrected by subtracting the product of ROI area by the mean fluorescence of multiple background ROIs measured per image. The Spc110-mCherry signal was used to localize the kinetochore. The dot plot graph displays individual corrected values. Horizontal bars represent the median value and error bars represent the 95% confidence interval. Significance was assessed with a two-tailed Mann-Whitney U test using GraphPad Prism (v10.6.1).

### Total internal reflection fluorescence microscopy (TIRFM) and analysis

TIRFM assay to detect de novo kinetochore assembly at single molecule resolution was performed as previously described (12, 23). Briefly, all images were captured using a Nikon TE-2000 inverted RING-TIRF microscope. Images were acquired at a resolution of 512 × 512 pixels with a pixel size of 0.11 μm/pixel at a readout speed of 10 MHz. Atto-647-labeled *CEN3* DNAs were excited at 640 nm for 300 ms, GFP-tagged proteins at 488 nm for 200 ms. Images were analyzed with CellProfiler (4.2.6) to assess colocalization and quantify signals between the DNA channel (647 nm) and GFP channel (488 nm). Results were processed and visualized using FIJI (ImageJ, v1.54p). The lysates were washed off after each time point before imaging. At least 3000 DNA molecules were imaged for each time point from each repeat. Error bars are standard deviation over three biological repeats.

### Analysis of gene expression

RNA from G1 arrested cells was extracted using TRIzol (Thermo Fisher, #15596) according to the manufacturer’s instructions. Contaminant genomic DNA was eliminated using TURBO DNA-free kit (Ambion, #AM1907). 1 μg of DNase-treated RNA was reverse transcribed using RevertAid Reverse Transcriptase (Thermo Fisher Scientific, #EP0442) in a 20 μL reaction using random hexamer and oligo(dT)18 primers and analyzed by qPCR using the Forget-Me-Not EvaGreen qPCR Master Mix (Biotium, #31045) with primers listed in SI appendix, Table S3. qPCR was performed using a Quantstudio™ 5 Real-Time PCR System (Applied Biosystem). A final dissociation stage was performed to verify the specificity of the PCR primers. Primer efficiency of each primer pair was evaluated previously (32). The relative expression of cenRNAs was normalized to *UBC6* expression, which is stably expressed throughout the cell cycle. The relative fold changes were calculated using the ΔΔCt method (60).

### Serial dilution assay

The designated *S. cerevisiae* strains were grown in YPD medium overnight. Cell concentration was measured on a spectrophotometer (Bio-Rad), and cells were diluted to OD600 = 1.0. Next, serial dilutions (1:5) were made in water in a 96-well plate and the wells were then spotted onto YPD or YC-methionine plates. Plates were grown for 3 days at 23 °C prior to imaging.

## Data availability

Mass spectrometry data generated in this study are available through Mass Spectrometry Interactive Virtual Environment (MassIVE, University of California San Diego) located here: https://massive.ucsd.edu/ProteoSAFe/dataset.jsp?task=22882ab4ea494f0288c94dfd7b2dc408.

## Acknowledgments

We thank Arshad Desai for the kind gift of antibodies and the Asbury and Biggins labs for insightful discussions. We thank the Biggins lab and Chip Asbury for feedback on the manuscript. We thank Mengqiu Jiang for her help with illustrations. Schematics in figures 1 and 3 were made using Biorender.com. This work was supported by NIH P30CA015704 that supports the Genomics and Bioinformatics Shared Resources of the Fred Hutchinson/University of Washington Cancer Consortium, NIH R35 GM149357/GM/NIGMS HHS/United States to S.B., and HHMI/Jane Coffin Childs Memorial Fund postdoctoral fellowship to C.H. S.B. is a Howard Hughes Medical Institute (HHMI) Investigator.

