## supplementary data file for "An interdependent Cbf1-CCAN interaction stabilizes the budding yeast kinetochore"

Sue Biggins

**This PDF file includes:**

Figures S1 to S4

Tables S1 to S3

SI References

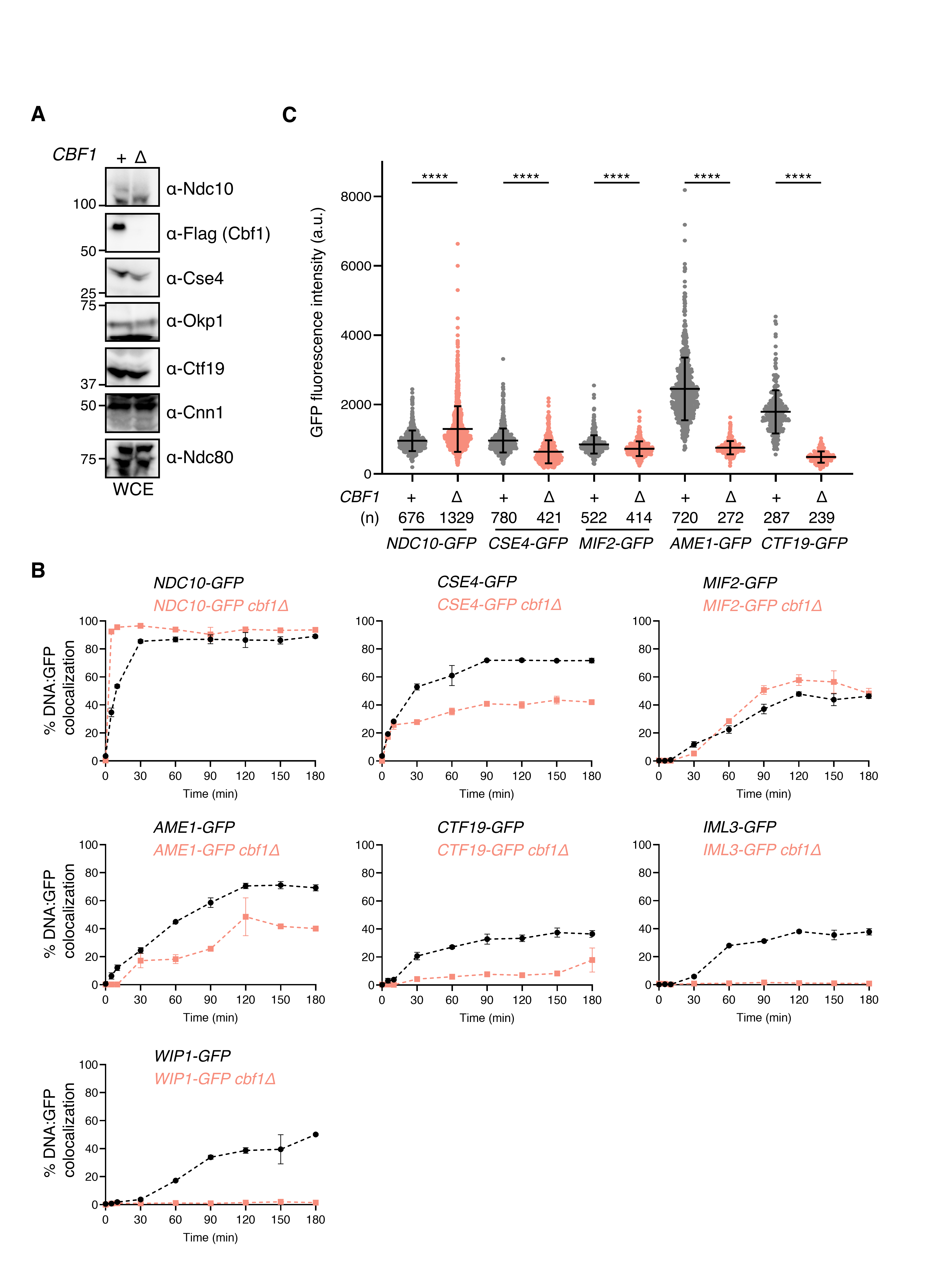
**Fig. S1. Cbf1 is required for kinetochore assembly in vitro.** (A) Whole cell extracts (WCE) from Fig. 1C were immunoblotted with the indicated antibodies. (B) Percentages of colocalization between *CEN3* DNAs and GFP-tagged protein in either in a wild type or *cbf1Δ* background as analyzed by TIRFM at various time points for 180 min. Strains used are as follows: *NDC10-GFP* (SBY22903), *NDC10-GFP cbf1Δ* (SBY24745*), CSE4-GFP* (SBY22195), *CSE4-GFP* *cbf1Δ* (SBY24743), *MIF2-GFP* (SBY22094), *MIF2-GFP* *cbf1Δ* (SBY24615), *AME1-GFP* (SBY22119), *AME1-GFP* *cbf1Δ* (SBY24617), *CTF19-GFP* (SBY22116), *CTF19-GFP* *cbf1Δ* (SBY24619), *IML3-GFP* (SBY22199), *IML3-GFP* *cbf1Δ* (SBY24621), *WIP1-GFP* (SBY22207), and *WIP1-GFP* *cbf1Δ* (SBY24625). Error bars represent the standard deviation over three biological repeats. At least 3000 DNA molecules were imaged for each biological replicate. Data for GFP-tagged proteins in a wild-type background were replotted from (1). (C) Fluorescence intensity analysis of GFP-tagged proteins after 180 min of kinetochore assembly. The horizontal lines represent the mean, and error bars represent the standard deviation. The number (n) of total kinetochore foci analyzed is noted on the graph. Statistical significance was assessed using a two-tailed Mann–Whitney U test (****, p<0.0001).

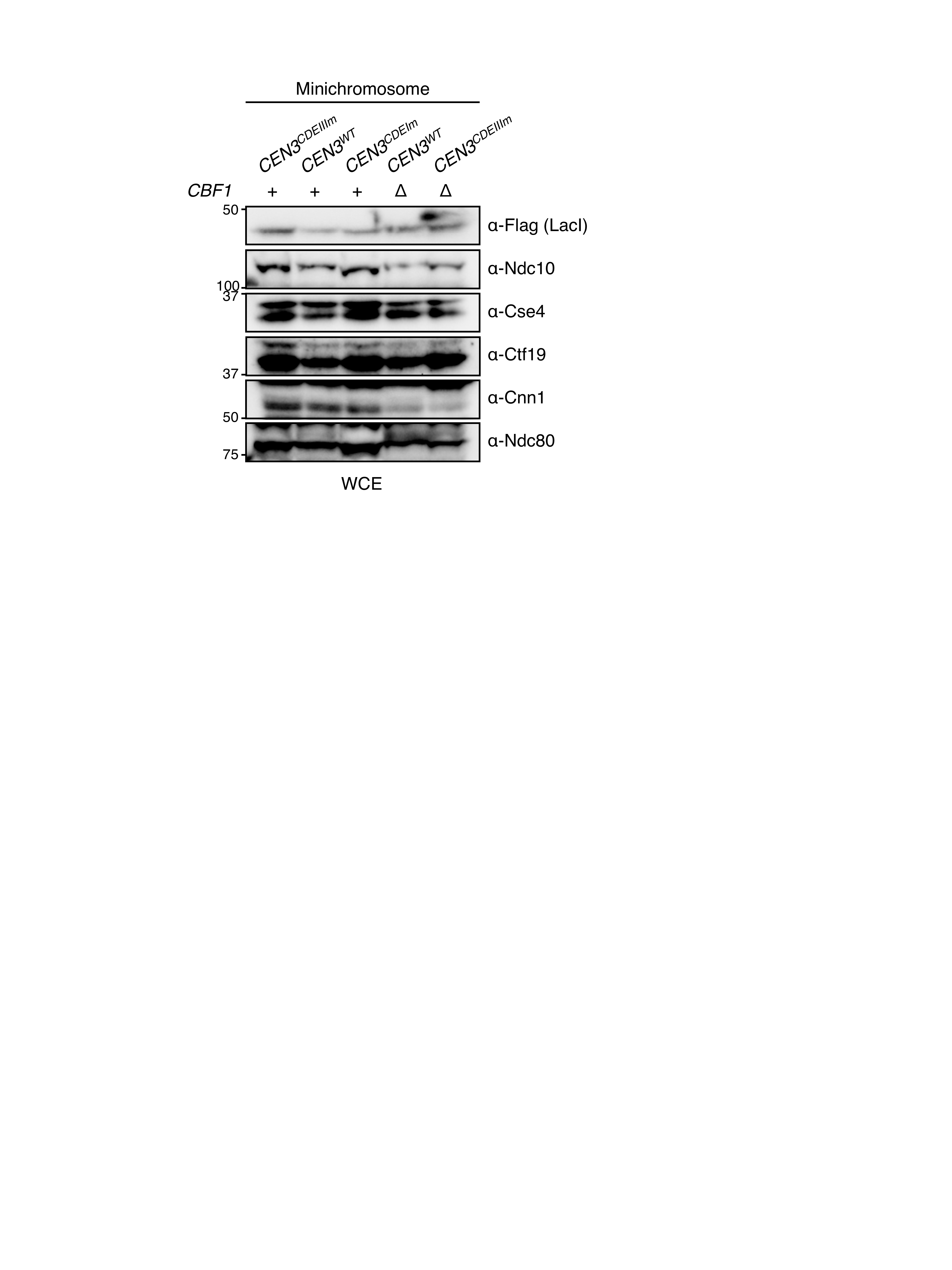
**Fig. S2. Cbf1 is required for kinetochore assembly in vivo.** Whole cell extracts (WCE) from Figure 2A were immunoblotted with the indicated antibodies.

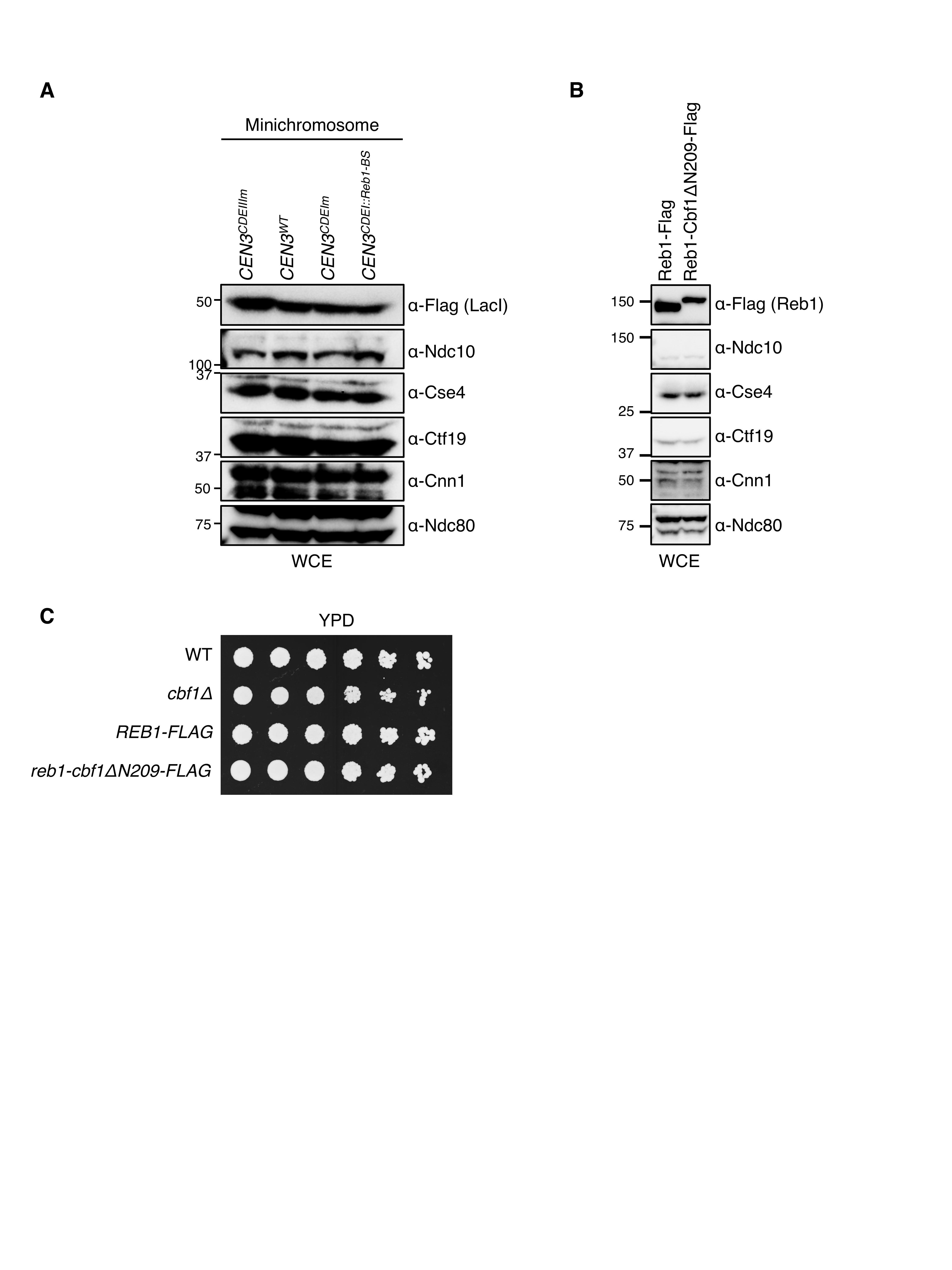
**Fig. S3.** **Cbf1’s role in CCAN recruitment is separate from its transcriptional roadblock activity.** (A) Whole cell extracts (WCE) from Figure 3B were immunoblotted with the indicated antibodies. (B) Whole cell extracts (WCE) from Figure 3D were immunoblotted with the indicated antibodies. (C) 5-fold serial dilution series of WT (SBY3), *cbf1Δ* (SBY4958), *REB1-FLAG* (SBY25215), and *reb1-cbf1ΔN209-FLAG* (SBY25217) strains were spotted on YPD and incubated for 3 days at 23 °C.

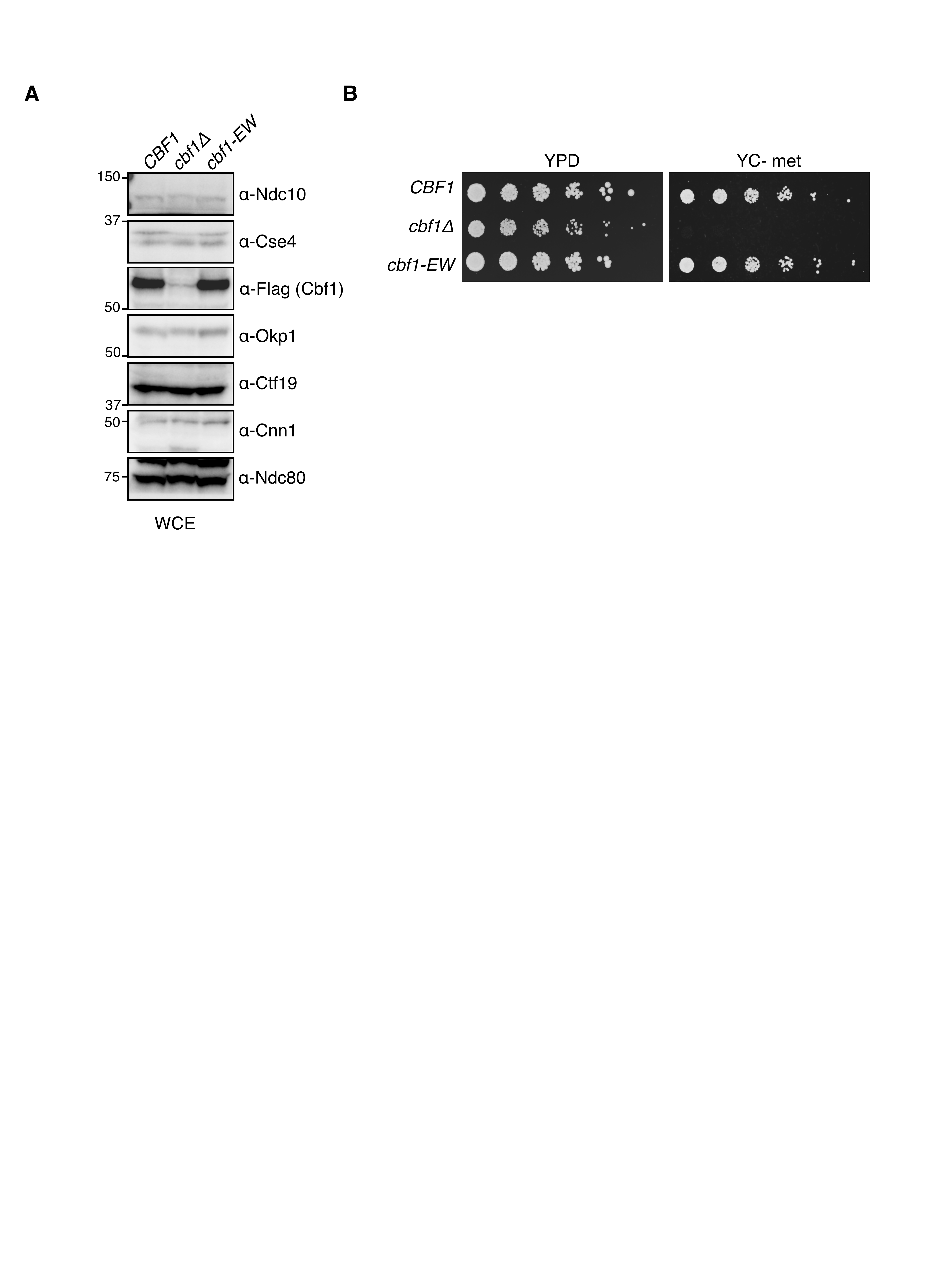
**Fig. S4. Cbf1-EW mutant disrupts CCAN assembly and derepresses CEN transcription.** (A) Whole cell extracts (WCE) from Figure 4C were immunoblotted with the indicated antibodies. (B) 5-fold serial dilution series of *CBF1-3FLAG* (SBY18421), *cbf1Δ* (SBY4958), and *cbf1-EW-3FLAG* (SBY22227) strains were spotted on YPD or YC-met (lacking methionine) plates and incubated for 3 days at 23 °C.

**Table S1. List of *S. cerevisiae* strains used in this study**

| **All strains are derivatives of SBY3 (W303)** | |  |
| --- | --- | --- |
| **Strain number** | **Genotype** | **Replicating plasmid** |
| SBY3 | *MATa ura3-1 leu2-3,112 his3-11 trp1-1 can1-100 ade2-1 bar1-1* |  |
| SBY18421 | *MATa CBF1-3xFLAG:TRP1* |  |
| SBY4958 | *MATa cbf1Δ::KanMX6* |  |
| SBY22903 | *MATα NDC10-GFP:KanMX6* |  |
| SBY24745 | *MATα NDC10-GFP:KanMX6 cbf1Δ::NAT* |  |
| SBY22195 | *MATα CSE4-GFP:NAT* |  |
| SBY24743 | *MATα CSE4-GFP:NAT cbf1Δ::KanMX6* |  |
| SBY22094 | *MATα MIF2-GFP:KanMX6* |  |
| SBY24615 | *MATα MIF2-GFP:KanMX6 cbf1Δ::NAT* |  |
| SBY22119 | *MATα AME1-GFP:KanMX6* |  |
| SBY24617 | *MATα AME1-GFP:KanMX6 cbf1Δ::NAT* |  |
| SBY22116 | *MATα CTF19-GFP:KanMX6* |  |
| SBY24619 | *MATα CTF19-GFP:KanMX6 cbf1Δ::NAT* |  |
| SBY22199 | *MATα IML3-GFP:KanMX6* |  |
| SBY24621 | *MATα IML3-GFP:KanMX6 cbf1Δ::NAT* |  |
| SBY22207 | *MATα WIP1-GFP:KanMX6* |  |
| SBY24625 | *MATα WIP1-GFP:KanMX6 cbf1Δ::NAT* |  |
| SBY19145 | *MATa ura3-1:pCMV-LACI-3FLAG:URA3 CNN1-3V5:KanMX CEN3-8LacO-TRP* | pSB963 |
| SBY19146 | *MATa ura3-1:pCMV-LACI-3FLAG:URA3 CNN1-3V5:KanMX CEN3^CDEIIIm^-8LacO-TRP* | pSB972 |
| SBY19147 | *MATa ura3-1:pCMV-LACI-3FLAG:URA3 CNN1-3V5:KanMX CEN3^CDEIm^-8LacO-TRP* | pSB2959 |
| SBY19148 | *MATa ura3-1:pCMV-LACI-3FLAG:URA3 CNN1-3V5:KanMX CEN3^CDEI::Reb1-BS^-8LacO-TRP* | pSB3118 |
| SBY19149 | *MATa ura3-1:pCMV-LACI-3FLAG:URA3 CNN1-3V5:KanMX CEN3-8LacO-TRP cbf1Δ::KanMX6* | pSB963 |
| SBY19150 | *MATa ura3-1:pCMV-LACI-3FLAG:URA3 CNN1-3V5:KanMX CEN3^CDEIIIm^-8LacO-TRP cbf1Δ::KanMX6* | pSB972 |
| SBY24879 | *MATa IML3-3mNeonGreen:HIS SPC110-mCherry::HPHMx* |  |
| SBY24881 | *MATa IML3-3mNeonGreen:HIS SPC110-mCherry::HPHMx cbf1Δ::KanMX6* |  |
| SBY14171 | *MATα dsn1-3A-3FLAG:URA3* |  |
| SBY22227 | *MATa cbf1-EW-3FLAG:TRP* |  |
| SBY22129 | *MATα CBF1-GFP:KanMX6* |  |
| SBY22923 | *MATα cbf1-EW-GFP:KanMX6* |  |
| SBY24889 | *MATα CBF1-GFP:KanMX6 ctf19Δ::KanMX6* |  |
| SBY22452 | *MATa CBF1-3xFLAG:TRP1 OKP1-3HA:HIS3* |  |
| SBY22454 | *MATa cbf1Δ::KanMX6 OKP1-3HA:HIS3* |  |
| SBY22456 | *MATa cbf1-EW-3xFLAG:TRP1 OKP1-3HA:HIS3* |  |
| SBY25315 | *MATa REB1-3FLAG:TRP1* |  |
| SBY25317 | *MATa REB1-cbf1ΔN209-3FLAG:TRP1* |  |

**Table S2. List of plasmids used in this study**

| **Plasmid name** | **Relevant genotype** | **Marker** | **Use** | **Origin** |
| --- | --- | --- | --- | --- |
| pSB963 | *CEN3* (WT) | Ampicillin, TRP | Kinetochore assembly assay (*CEN3* WT and *ampC* templates) | (2) |
| pSB972 | *cen3* (CDEIIIm) | Ampicillin, TRP | Kinetochore assembly assay (*CDEIIIm* template) | (2) |
| pSB2959 | *cen3* (CDEIm) | Ampicillin, TRP | Kinetochore assembly assay (*CDEIm* template) | (3) |
| pSB3118 | *cen3* (CDEI::Reb1-BS) | Ampicillin, TRP | Kinetochore assembly assay (*CDEI::Reb1-BS* template) | (3) |
| pSB3006 | *CBF1-3FLAG:TRP* | Ampicillin | Template for fusing *cbf1-ΔN209-3FLAG:TRP* to *REB1* at its C-terminus | This study |
| pSB3744 | *cbf1-EW-3FLAG:TRP* | Ampicillin | For replacement of endogenous *CBF1* locus with *cbf1-EW-3FLAG:TRP* | This study |

**Table S3. List of primers used in this study**

| **Target** | **Primer name** | **DNA sequence (5' to 3')** | **Application** | **Comment / Purpose** |
| --- | --- | --- | --- | --- |
| CEN3 | SB3878 | BIOTIN-GGTTCTGGTGGTTCTGGTGGTTCTGGTGAATTCAAACAACCGCCGGCTTCCACCA | PCR | Amplification of CEN3 DNA from plasmids pSB963, pSB972, pSB2959, or pSB3118 to be used as template for the kinetochore assembly assay |
|  | SB3880 | ATCAGCGCCAAACAATATGGAA |  |  |
| AME1 | SB9009 | GAAAAAAATAAATAAAATTAATGAAAATCTTTCTAACGAATTACAACCAAGTCTAGGTGACGGTGCTGGTTTA | PCR | Deletion of *CBF1* locus using pFA6a plasmids |
|  | SB9010 | AATACATATATACATATATATATATATATATATATACATCTTTTGAACCAATTCCTCGATGAATTCGAGCTCG |  |  |
| CBF1 | SB1058 | CAACATCAAGTGCTTAAAATATAATACGGTTTTCTACACTTTTATTAACGCGGATCCCCGGGTTAATTAA | PCR | To tag *AME1* in C-ter using pFA6a plasmids |
|  | SB1059 | GCAGATACATAGGGAGACTCGAAATACATTTAGCTATCTATTTTTAACTCTCGATGAATTCGAGCTCGTT |  |  |
| CBF1 | SB5550 | CGAAAGAAAAAGCACTAGGAGCGATAATCCACATGAGGCTAGGGAACAAAAGCTGGAGCT | PCR | To tag *CBF1* in C-ter using pFA6a plasmids |
|  | SB5551 | AGGGAGACTCGAAATACATTTAGCTATCTATTTTTAACTCCTATAGGGCGAATTGGGT |  |  |
| CNN1 | SB5152 | TCCCTTAGAACTTCAATCAAGAATTGAAAGTTATTTGTTCCGGATCCCCGGGTTAATTAA | PCR | To tag *CNN1* in C-ter using pFA6a plasmids |
|  | SB5153 | AAATTGCCACTATTTAATTATTTTCTCTACGGTATCTTTTGAATTCGAGCTCGTTTAAAC |  |  |
| CTF19 | SB9027 | AACCGGGTTAAAGGAGATCTGCAACGTTTGCCTATTCCCGGACATGTACGCCAGGGGTGACGGTGCTGGTTTA | PCR | To tag *CTF19* in C-ter using pFA6a plasmids |
|  | SB9028 | GAGCTTATCGGAATCGTTTAAGCAAGCCGTCCAGTTGGCAATGGCAAATGGAACATCGATGAATTCGAGCTCG |  |  |
| IML3 | SB9043 | TCTCAGGAACAGAGTTCTAGCAGTTGTACTCCAATCGATTCAGTTTACCAGCGAGGGTGACGGTGCTGGTTTA | PCR | To tag *IML3* in C-ter using pFA6a plasmids |
|  | SB9044 | AAAAAAGGTAGAGCTGTGGTTTTTTATTGTATCTTGGTGAATATTCTTTATAGTGTCGATGAATTCGAGCTCG |  |  |
| MIF2 | SB9015 | AGACGCTAACGATGACAACGACAAAGAATTAGACAGTACGTTTGACACTTTTGGGGGTGACGGTGCTGGTTTA | PCR | To tag *MIF2* in C-ter using pFA6a plasmids |
|  | SB9016 | CCTAGTTATATTTCTTCAGTACATAGCATGCATAATGAGAATATTCACATCATAATCGATGAATTCGAGCTCG |  |  |
| NDC10 | SB9041 | GTGGAGGCATGACCATCAAAATTCATTTGATGGTCTGTTAGTATATCTATCTAACGGTGACGGTGCTGGTTTA | PCR | To tag *NDC10* in C-ter using pFA6a plasmids |
|  | SB9042 | TATAAACATACATGTCGGTATCCCTATACGAAACAGTTTAAACTTCGAAGCTCCCTCGATGAATTCGAGCTCG |  |  |
| REB1 | SB6222 | TGATTATTTTAGCTCCAATATTTCAATGAAAACAGAAAATCGGATCCCCGGGTTAATTAA | PCR | To tag *REB1* in C-ter using pFA6a plasmids |
|  | SB6223 | TTATTGAGTTTTTCGCTTTCACCAATTATATTTTCCGGAAGAATTCGAGCTCGTTTAAAC |  |  |
| REB1 | SB6266 | TGATTATTTTAGCTCCAATATTTCAATGAAAACAGAAAATCCTACTACTTTGGCCACAAC | PCR | To tag *REB1* in C-ter with *cbf1-ΔN209-3FLAG:TRP* |
|  | SB6267 | TTATTGAGTTTTTCGCTTTCACCAATTATATTTTCCGGAAGGCAAGTGCACAAACAATAC |  |  |
| WIP1 | SB9039 | GGAAGTATATCCTTATCATATTGAAGCTGCAACGCAGGCTTTTCTGGATAGTCAAGGTGACGGTGCTGGTTTA | PCR | To tag *WIP1* in C-ter using pFA6a plasmids |
|  | SB9040 | TTATTTGCTATTACGAACAAAAGAGTATATGATAAAGAGGCTTAAAAATACCCCTTCGATGAATTCGAGCTCG |  |  |
| CBF1 | SB5798 | ATTTAAATTTATGCTTTAGTATCGTCATATTC | Cloning | To clone *CBF1-3FLAG:TRP* into a pCRII-TOPO vector |
|  | SB5799 | ATTTAAATAAAGACATATTTGAAAGTCCGTC |  |  |
| CBF1 | SB7917 | ATTGTGGAGCGAGCAAAACGCATCG | Cloning | Site-directed mutagenesis of L283E and L287W on plasmid pSB3006 |
|  | SB7918 | TTTTGTTCCGTCCACTTTTCGATGTTTGC |  |  |
| CEN8 | SB7232 | AACTCCAACAATTACACATCCACAAAACG | RT-qPCR | To amplify cenRNA from *CEN8* |
|  | SB7233 | TTTCTAAGTTCGGAACACAAAACCCAATG |  |  |
| CEN4 | SB6685 | GATTACCGAAACATAAAACCTGCTCAAG | RT-qPCR | To amplify cenRNA from *CEN4* |
|  | SB6795 | TATGAAAGCCTCGGCATTTTGGC |  |  |
| CEN5 | SB6721 | AGCAGTATTAGATTTCCGAAAAGAAAAAAAGG | RT-qPCR | To amplify cenRNA from *CEN5* |
|  | SB6457 | ACAACTGCTATTTATGTGCGGC |  |  |
| UBC6 | SB4137 | GATACTTGGAATCCTGGCTGGTCTGTCTC | RT-qPCR | To amplify UBC6 transcript to be used as a RT-qPCR normalizer |
|  | SB4138 | AAAGGGTCTTCTGTTTCATCACCTGTATTTGC |  |  |
